# FastProject: A Tool for Low-Dimensional Analysis of Single-Cell RNA-Seq Data

**DOI:** 10.1101/043463

**Authors:** David DeTomaso, Nir Yosef

## Abstract

**Background:** A key challenge in the emerging field of single-cell RNA-Seq is to characterize phenotypic diversity between cells and visualize this information in an informative manner. A common technique when dealing with high-dimensional data is to project the data to 2 or 3 dimensions for visualization. However, there are a variety of methods to achieve this result and once projected, it can be difficult to ascribe biological significance to the observed features. Additionally, when analyzing single-cell data, the relationship between cells can be obscured by technical confounders such as variable gene capture rates.

**Results:** To aid in the analysis and interpretation of single-cell RNA-Seq data, we have developed FastProject, a software tool which analyzes a gene expression matrix and produces a dynamic output report in which two-dimensional projections of the data can be explored. Annotated gene sets (referred to as gene ‘signatures’) are incorporated so that features in the projections can be understood in relation to the biological processes they might represent. FastProject provides a novel method of scoring each cell against a gene signature so as to minimize the effect of missed transcripts as well as a method to rank signature-projection pairings so that meaningful associations can be quickly identified. Additionally, FastProject is written with a modular architecture and designed to serve as a platform for incorporating and comparing new projection methods and gene selection algorithms.

**Conclusions:** Here we present FastProject, a software package for two-dimensional visualization of single cell data, which utilizes a plethora of projection methods and provides a way to systematically investigate the biological relevance of these low dimensional representations by incorporating domain knowledge.

## Background

In an analogous manner to the maturation of RNA-Seq methodologies, single-cell RNA-Seq (scRNA-Seq) is now in its infancy and requires new computational tools to realize its full potential for dissecting and understanding the functional meaning of cell-to-cell heterogeneity [1, 2]. Visualization methods provide an effective strategy for inspecting and characterizing the phenotypic diversity between cells. In a typical scenario, the analysis begins with a matrix of expression levels of thousands of genes in hundreds of cells. An appealing way to make sense out of this immense data is to project it into a two dimensional scatter-plot, where each cell is represented by a single point. While such representations provide an easy way to see obvious stratification of cells into a taxonomy of discrete types, they can also provide more nuanced views of gradual transitions, reflecting for instance, developmental processes [3], physical locations [4], or the cell cycle [5]. Indeed, supplementing these two-dimensional views with additional, phenotypical information (e.g. the expression level of marker genes) can be used to provide the correct context, and make the observed diversity between cells interpretable [6, 7]).

There are three main challenges in making effective use of such visualizations for scRNA-Seq data. The first challenge is selecting an appropriate method for dimensionality reduction (projection) among candidates such as principal component analysis (PCA) [1, 2], independent component analysis (ICA) [3] or various non-linear methods such as t-distributed stochastic neighbor embedding (t-SNE) [8], each of which may highlight different aspects of the data. Once a projection is created, a second challenge is to interpret its biological significance, namely which cellular phenotypes [7] or processes [9] are most responsible for the resulting arrangement of cells. Lastly, scRNA-Seq can be difficult to correctly interpret due to technical confounders such as differences in gene capture rates. Performing functional interpretation on the input data without accounting for this effect may lead to incorrect interpretation of the biological meaning of cell-to-cell heterogeneity.

## Introducing FastProject

To address these issues, we have developed FastProject, a software tool for the visualization and interpretation of scRNA-Seq data. FastProject allows the user to explore a gene expression matrix using a plethora of two-dimensional visualization methods. To interpret these two-dimensional plots, we use the concept of biological signatures - sets of genes that represent a dichotomy between two cellular states of interest [7] (e.g. epithelial to mesenchymal transition [2]). We evaluate the extent to which these phenotypic dichotomies are reflected in the projections (i.e. to what extent do neighboring cells in the projection have similar values for the genes included in the signature), and highlight the significant projection-signature pairs. This analysis is made possible in single cells by modeling the probability that a missed transcript was actually expressed in the cell, and using these probabilities when evaluating signature scores on samples to minimize the effect of variable capture rates between cells. As a source for biological signatures, we use publicly available datasets that consist of comparative information from hundreds of studies (e.g. MSigDB [10]), which can be supplemented by the user to include any other gene signatures of interest. Through this automated analysis, FastProject therefore provides a systematic view of the main axes of variation in the data, along with their possible biological meaning.

It is important to clarify up-front that FastProject is not a normalization tool. Indeed, it has been observed by us and others that without proper normalization scRNA-Seq data can be heavily confounded by technical factors such as library depth and complexity [7]. We strongly advocate the use of scRNA-Seq normalization tools (e.g. based on [5] or [7]) prior to any downstream analysis, and we assume that the data has been normalized prior to input to FastProject. Nevertheless, since scRNA-Seq tend to be characterized by strong zero-inflation, we conduct our biological signatures analysis while paying close attention and controlling for possible effects of gene dropout events (false negatives).

When running FastProject, processing is done upfront (10-30 minutes on typical data sets), producing dozens of projection maps (using different algorithms and parameterizations) and their functional annotation, which can be inspected through an interactive HTML report. Because processing is performed upfront, the user can quickly switch between different projection maps in the output report as well as share the results with colleagues who would not themselves need to install FastProject. Importantly, FastProject has been written to be easily extensible so that it may serve as a general platform for deploying and evaluating new gene filtering schemes, false-negative estimation methods, or projection techniques. Instructions for developers on how to augment FastProject are detailed in the FastProject wiki hosted at https://github.com/YosefLab/FastProject/wiki.

## Implementation

### Overview

The steps involved in the FastProject processing pipeline are depicted in Figure 1. The software starts with an evaluation of false negative rates, later used to down-weight the effect of drop-outs on the biological signature analysis. It then selects sufficiently variable genes for further analysis using increasingly stringent criteria. With these genes in hand, FastProject uses 11 different projection methods (summarized below) to calculate two dimensional coordinate for each cell. Based on a user-provided database of gene signatures it then computes a score for every cell/signature pair and uses a randomization test to identify statistically significant projection-signature associations. Importantly, the confounding effect of missed transcripts is mitigated by estimating the probability that each undetected gene was actually expressed in the cell, and down-weighting the contributions of these measurements in downstream analysis (similar to [7]). Altogether, FastProject outputs 76 possible projec-tions (a combination of choosing different gene filtering criteria, whether or not the data was reduced to significant PCs prior to projection, and the final projection method) along with their functional annotations, which can be interactively inspected through a user-friendly HTML report (Figures 1b, and 3). The results are also provided in the form of text files (including signature scores, projection coordinates etc.), which can be used for downstream analysis.

**Figure 1:**
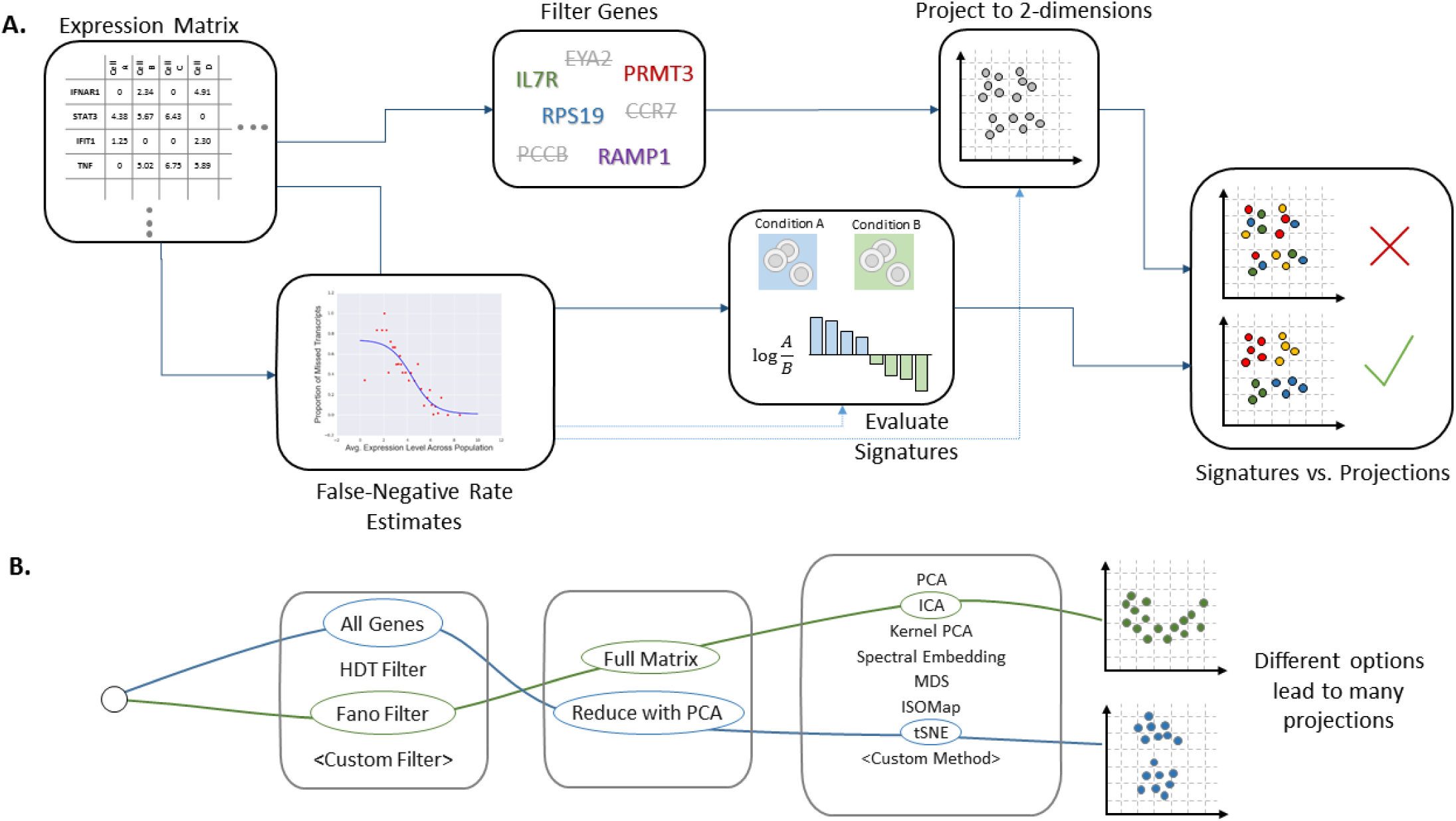
The FastProject Pipeline. A) Diagram describing the FastProject pipeline. A gene expression matrix is taken as input (left), and the resulting output report (right) combines low dimensional-representations of the input with gene signatures to highlight signatures which best explain features in the data. B) Configurations for the projection that can be selected among in the output report.

**Figure 2:**
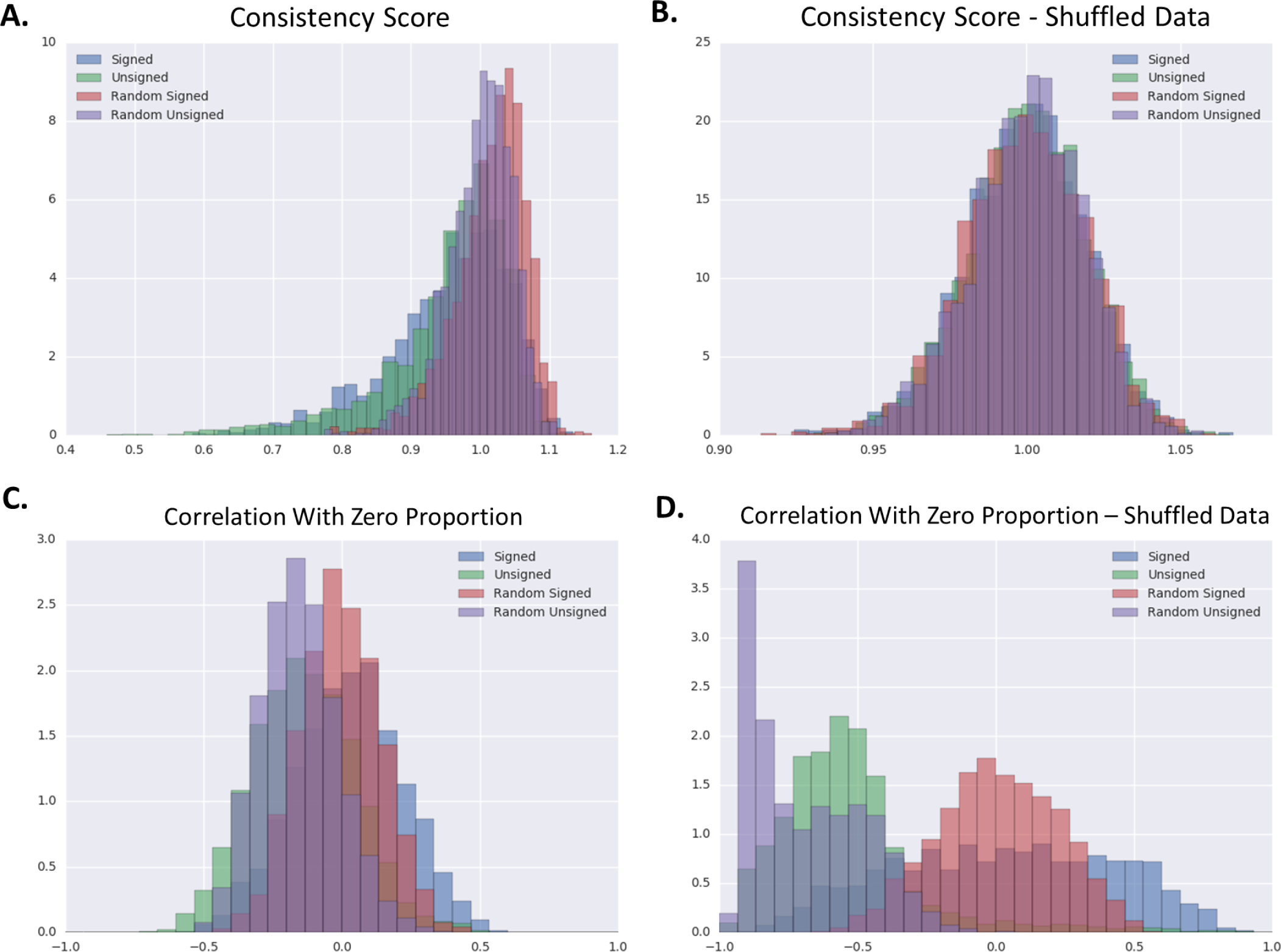
Behavior of Signature Scores. Behavior of signature scores calculated on the human glioblastoma scRNA-seq data from Patel et al, 2014 [2]. A) Distribution of Signature/Projection consistency scores across four different types of signatures, Signed (signed immunological signatures from MSigDB), Unsigned (various unsigned hallmark and pathway signatures from MSigDB), Random Signed (signed signatures with randomly selected genes), and Random Unsigned (unsigned signatures with randomly selected genes). Consistency scores normalized by the mean of the Random Unsigned scores. B) Distribution of Signature/Projection consistency scores for data in which gene expression levels have been shuffled within each cell. C) Distribution of the Pearson’s correlation coefficient between signature scores and a confounding variable - the proportion of undetected genes in a sample. D) Same as C), but signature scores have been calculated by simply taking the unweighted average of log expression level for genes in the signature. Note how with the simple method in D, signatures tend to be much more strongly correlated with the confounding variable.

**Figure 3:**
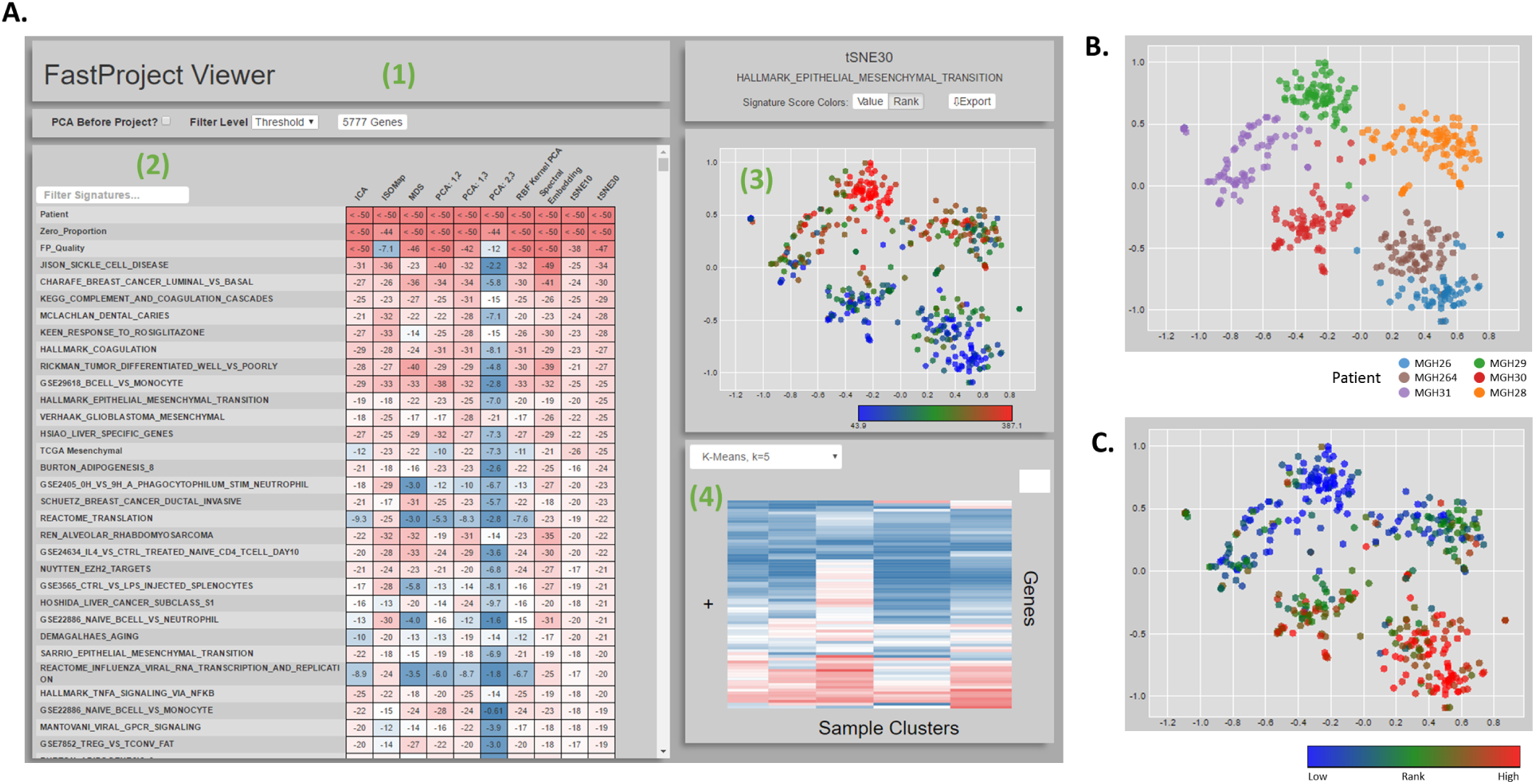
FastProject Output Report. Screenshot of FastProject interactive output report. 1) Controls for changing which genes were used when generating the projection and whether or not PCA was applied first. 2) Table displaying significance of the consistency score for each signature/projection pairing. Each row represents a signature and each column, a projection method. Clicking a cell in the table selects a signature and projection. 3) Scatter plot showing the selected projection annotated (color) with signature scores from the selected signature. 4) Heatmap showing average expression level of genes within each cluster. The clustering method can be changed through the dropdown menu in the same panel. B) Corresponding scatterplot when selecting projection tSNE30 and the Patient signature. C) Scatterplot for a signature representing response to the PPAR_*γ*_ agonist rosiglitazone.

### False-Negative estimates

To account for expressed transcripts that are not detected (false negatives) due to the limitations in sensitivity [1, 11], an initial step in the processing pipeline is to evaluate these rates so that the subsequent analysis can down-weight the contribution of less reliably measured transcripts. This is done by estimating a false negative rate for each cell individually (as different cells within a sample can have different levels of library quality and cell integrity) using a set of human housekeeping genes [12]. Our procedure derives from the assumption that housekeeping genes are missed due to technical errors (i.e. all housekeeping genes are constitutively expressed), and that the probability of missing a transcript is related to its average expression level in the expressing cells. Importantly, as the appropriate set of constitutively expressed genes may differ from study to study and between organisms, FastProject can accept a user-defined housekeeping list. Our estimation of false negative rates is built on and extends upon our previous work [7]. For each housekeeping gene, we estimate its mean expression level by taking the average of non-zero measurements for the gene. We then use the estimated means to group the genes into 30 quantiles, and denote the mean of genes in quantile 1 ≤ *q* ≤ 30 as *μ*_*q*_. For each cell *j* and quantile q, we then compute *F*_*qj*_ as the proportion of genes from q that are not detected by *j*. Based on our assumption of constitutive expression, we treat *F*_*iq*_ as an empirical estimate to the dropout rates (i.e. probability that a gene is not observed, conditioned that it is expressed). We use the *F*_*jq*_ values to fit a sigmoid function *F*_j_(·) that describes the observed dropout rates as a function the genes’ average expression when detected (Figure 1A):

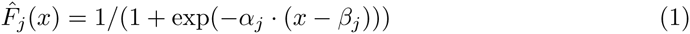

where the estimated parameters *α*_*j*_ and *β*_j_ minimize the residual sum of squares (RSS) term:

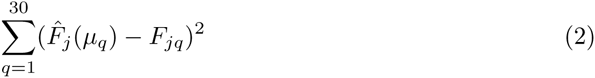

Applying the fitted function globally for all genes, we estimate the conditional probability for a dropout event for gene *i* in cell *j* as: *P*(Gene *i* is not detected in cell *j* | Gene *i* is expressed by cell j) = *F̂*_*j*_(*μ*_*i*_) where *μ*_*i*_ is the average of gene *i*’s expression level when detected. Finally, we estimate the prior probabilities for detection and expression of each gene in order to evaluate the false negative probabilities, *P*(Gene i is expressed by cell j | Gene i is not detected in cell j), as described in the Methods section. These probability estimates are used further in the pipeline to reduce the effect of missed transcripts when evaluating signature scores and generating projections.

The fitted sigmoids *F̂*_*j*_(·) can also be used to provide an overall evaluation of the abundance of false negatives in each cell *j* by taking the area under the curve. This in turn provides a way to identify and exclude poor quality samples, assuming that such samples have higher dropout rates. Such quality control filter (on cells) is available in FastProject and can be enabled when running the pipeline (it is turned off by default). With this option enabled, samples that score 1.6 median absolute deviation (MAD) units worse than the median quality score are removed prior to calculating signatures and low-dimensional projections.

## Generating 2-dimensional projections

### Gene filtering

Selecting an informative set of genes is necessary for obtaining biologically meaningful patterns of variability between cells. To this end, FastProject applies a strict pre-filtering step that discards genes detected in less than a threshold number of cells. The default threshold is 20% of the input cells, however this is configurable as an input option. Subsequently, FastProject produces projections that derive from all pre-filtered genes as well as subsets thereof, calculated using two filtering schemes. The first selects bi-modal genes, using the Hartigan’s Dip Test (*p* < 0.05). The second selects highly-variable genes, based on their Fano-factor (*σ*^2^/*μ* where *μ* is the average expression and σ the standard deviation across all cells). To select candidates with high Fano-factor, genes are first stratified into 30 quantiles according to *μ*, and within each quantile genes are retained if their Fano-factor is more than 2 MAD above the quantile’s median Fano-factor.

### Projection Methods

The variety of methods available for the task of dimensionality reduction each come with strengths and weaknesses. For instance, projections of scRNA-Seq data based on PCA [7], provide an appealing guarantee about the preservation of variation, and makes the contribution of individual genes readily interpretable [6]. However, the underlying assumptions of PCA may not necessarily be supported by single cell data. In particular, PCA is a linear transformation, which may not be able to accurately capture non-linear relationships in the data (i.e. if the data is embedded within a non-linear low-dimensional manifold). Additionally, PCA posits that the projection axes should be uncorrelated, which again may not necessarily result in the most informative representation. The same criticisms apply for other linear methods such as ICA[3]. A complementary, and commonly used approach (t-SNE[8]) uses a non-linear projection that aims to preserve the structure in the data locally. However, while this method aims to ensure that points that are close in the high dimensional space will be close (with high probability) in the low dimensionality embedding, more global relations are not directly interpretable from the results. To avoid the issue of choosing up-front between these different options, FastProject uses these methods plus additional non-linear projection methods, including: ISOMAP [13] (using four nearest neighbors when defining the adjacency graph), PCA with a radial basis function (RBF) kernel (with kernel coefficient of 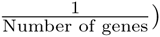, Multi-Dimensional Scaling (MDS), and Spectral Embedding [14](with Graph laplacian formed using k-nearest neighbors with 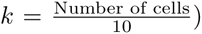. For the linear PCA we consider three combinations of principal components (1st and 2nd, 1st and 3rd, 2nd and 3rd); and for tSNE we use two perplexity values (10 and 30), totaling in 11 projection methods overall. All methods are run as implemented in the Scikit-Learn package for Python[15]. After each projection, the resulting sets of 2-dimensional coordinates are mean-centered and scaled such that the average *r*^2^ = [*x coordinate*]^2^ + [*y coordinate*]^2^ equals 1. This standardization is performed so that signature-projection scores (defined below) are more comparable between projections.

### Incorporating False-Negative estimates into Projections

Non-linear methods such as t-SNE have been shown to effectively combine with PCA. In this approach, PCA is performed first (using only highly variable genes), and only PCs that explain significantly more variance than expected by chance are retained. The resulting low-dimensional data points (yet of dimension of typically more than 2), are postulated to represent a cleaner version of the data, and are then embedded in two dimensions using the non-linear approaches. To allow for this option, FastProject generates two outputs for each non-linear projection method: with and without PCA prior to projection. To accomplish this, the software employs a randomization scheme similar to [16] to select the top principal components with statistically significant contribution to the overall variance (*p* < 0.05). The number of PCs selected by this procedure is enforced to be at least 5. All the subsequent non-linear projections are done based on this reduced-dimension matrix (in a typical scRNA-Seq dataset, the selected number of PCs is between 10 to 15). To provide a way for evaluating the effects of this reduction, FastProject also runs the complete analysis on the original, non-reduced matrix. When PCA is performed (either as a preceding step to the non-linear projections, or as a separate projection method), we use a weighted covariance matrix to account for the false negative rate estimates (similar to [7]). Entries in the weighted covariance matrix are calculated as:

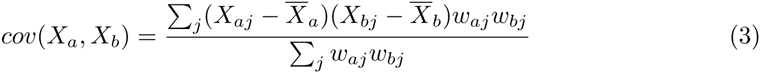

where X_*ij*_ is the log-transformed expression of gene *i* in cell *j* and *w* represents the matrix of weights of equivalent size, defined as:

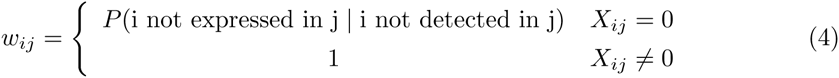

### Signature-Based Analysis

To interpret the biological meaning of the organization of cells in the resulting two dimensional maps, FastProject incorporates domain knowledge through the input of gene “signatures” [7]. The signatures can reflect a comparative analysis between two conditions of interest and consist of a set of genes, each of which is labeled as either ”up-regulated” or “down-regulated” in that comparison. Signatures, such as these, are based off of annotations of cell states obtained from public databases (e.g. the Immsig collection[10]), or provided by the user. For each signature, a score is computed against each cell by aggregating over the weighted expression level of its genes. Specifically, for signature *S* and cell *j*, the respective signature score *R*_*s*_(*j*) is calculated as:
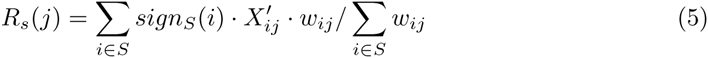

Where 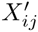 is the standardized (Z-normalized across all cells) log expression level of gene *i* in cell *j*, *W*_*ij*_ is the estimated False-Negative weight (defined above), and *signs*(*i*) = —1 for genes in the “down-regulated” set and +1 otherwise. Notably, signatures can also be undirected, in which case the sign value is set to +1 for all member genes.

### Projections vs. Signatures

A signature-projection consistency score is calculated to evaluate how well each projection reflects the phenotypic variation that is captured by each signature. To this end, for each pair of signature, *s*, and projection, *p*, we compute a signature consistency score representing the extent to which neighboring points in the projection have similar signature scores. This is done by calculating for each cell *j*, an estimated signature rank *r̂*_*sp*_(*j*) based on its location in the two dimensional plot:

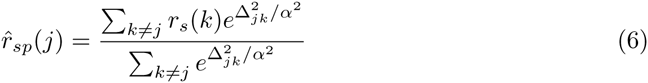

where Δ_*jk*_ is the Euclidean distance between cells *j* and *k* in the projection, *α* defines an effective neighborhood size (set to 0.33 by default), and *r*_*s*_(*j*) is the rank of the signature score for cell *j* (i.e. a rank transformation of the quantitative signature scores *R*_*S*_). The signature-projection consistency is then determined by the respective goodness of fit:

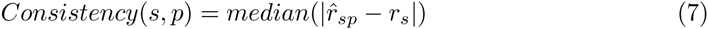

In this way, each signature/projection pairing is scored based on how similar signature scores are for samples located nearby in the projection space. To identify significantly consistent signature/projection pairs, we use signatures of randomly selected genes to create empirical background distributions of signature scores. We compare the consistency scores computed for the original signatures with those of the random ones, obtaining P-values using a Z-test and correcting for multiple hypothesis testing using the Benjamini Hochberg procedure. We observed that the distribution parameters of scores generated by random signatures varied with the number of genes in the signature. To account for this, separate distributions are generated for different signature sizes (10, 20, 50, 100, and 200 genes) and when assessing the significant of a signature score, the score is compared against the background distribution with the most similar number of genes. In the output, we report all the signatures that had a significant match (*FDR* < 0.05) with at least one projection.

## Results

### Software

FastProject has been implemented as a command line Python package. As an input the software receives: (1) an expression matrix in a tab-delimited format (genes in rows, cells in columns). (2) a set of gene signatures, using the standard GMT format. Such sets of directed signatures are publicly available from various databases, such as MsigDB[10] (e.g.including signaling effects of genetic and chemical perturbations, cell cycle signatures, and comparison of cell types) and NetPath[17] (transcriptional effects of signaling cascades). Un-directed gene sets are naturally more abundant, and can be drawn from resources such as Gene Ontology[18], KEGG[19, 20], and MSigDB[10] (note that the latter includes much of the information in the former two). Importantly, the user can also upload his/her own signatures that reflect a phenotype of interest. For instance, in the example below, we study glioblastoma cells, and use signatures derived from microarray experiments, which define different tumor sub-types. FastProject also accepts pre-computed signatures, namely, an association of cells with pre-computed values. These can be categorical (e.g. annotations of clusters, computed by a different tool); or numerical (e.g. additional meta data associated with cells; e.g. levels of a handful of proteins, or technical factors such as library complexity; see Methods section for description of how pre-computed signatures are analyzed).

FastProject will calculate projections, signature scores, and their associations, covering all the options above (totaling in 76 different projections; Figure 1b). To examine the extent to which the projections are affected by zero values, FastProject treats the sample quality score (defined above), and the percentage of genes with zero expression as pre-computed signatures and evaluates their association with each projection method. The output is provided as an HTML file (Figure 3), where projections, signatures and their associations can be inspected interactively. Additionally, a data export feature embedded in the HTML report allows the generation of tab-delimited text files that depict the output projection coordinates, signature scores, and other relevant information. The source code, running, examples and user manual are bundled with this manuscript submission and also hosted at https://github.com/YosefLab/FastProject.

### Extending FastProject

FastProject has been designed using a modular architecture so that new projection methods or criteria to filter genes can easily be added to the pipeline. Since dimensionality reduction is still an active research area, this allows new methods to easily be compared against more traditional approaches. For example, the recently proposed ZIFA algorithm[21] can be added by appending the following lines to Projections.py:

~~~
from ZIFA import block_ZIFA , ZIFA
def apply_ZIFA(proj_data, proj_weights=None):
   Z , model params = ZIFA. fitModel (projdata .T, 2) ;
   return Z;
_proj_methods [ ’ZIFA ’ ] = apply_ZIFA ;
~~~

This is documented in the software wiki, hosted with the project repository at https://github.com/YosefLab/FastProject/wiki.

### Proof of Concept

We applied FastProject to a recently published data sets of tumor cells from five glioblastoma patients[2]. The analyzed data, consisting of 430 single cells with mRNA abundance in units of transcripts per million (TPM) and normalized as described in [2], was downloaded from the Gene Expression Omnibus, accession number GSE57872. We applied FastProject on this data, using a large collection of both ’’signed” (i.e. up‐ and down regulated genes) and ’’unsigned” signatures from MSig DB (the C2 (curated), H (Hallmark), and C7 (Immune signatures) collections) and NetPath[17]. As a first check, we explored the distributions of signature consistency scores obtained for the original vs. random signatures, and compared the results to an application of FastProject on randomized datasets, where the entries in each column (Cell) were shuffled (i.e. maintaining the percentage of zeroes in every cell; Figures 2a-b). As expected, we see pronounced differences between the original input signatures and the randomized ones when FastProject is applied on the non-shuffled data, and these differences disappear when we apply FastProject on the randomized data. As a second test, we evaluated the extent by which the signature consistency scores are biased by the abundance of zero-values in the analyzed cells. As expected, when we do not down-weigh the potential false negative entries, the signature consistency scores highly correlate with the amount of zeroes in each cell; namely the analysis primarily reflect what might be a result of technical dropouts. Conversely, down-weighing the false negatives corrects this bias (Figures 2c-d). We repeated this procedure using a second dataset of scRNA-Seq data from mouse dendritic cells responding to antigen stimulation [1], obtaining similar results (Figure S1).

Examining the output report of FastProject (Figure 3a), we first observe that the various projection methods correctly stratify the cells according to their respective donors (Figure 3b). More importantly, FastProject automatically picks up on several of the most important features in this data, providing a proof of concept for its utility as an unbiased analysis tool. Specifically, the two dimensional position of the cells is highly consistent with their scoring according to an epithelial to mesenchymal transition signature, which is a strong marker of poor prognosis[22, 23]. The two dimensional positions are also associated with signatures of other key pathways altered in cancer, including immune and hypoxic responses as detailed below. While these observations were made using a general database of signatures (MSigDB), we supplemented our analysis with case-specific signatures - in this case gene signatures from TCGA that are predictive of Glioblastoma subtypes[23] to provide further support. As expected we see high level of concordance between the TCGA-derived scores of the Mesenchymal tumor sub-type and the epithelial to mesenchymal transition signatures from MSigDB. We also see the mirror image of the cell ranking when we consider the TCGA-derived signature of the Proneural tumor subtype of glioblastoma.

In addition to the HTML report, FastProject outputs all the cell-signature scores in textual format. Taking advantage of this feature, we were able to more closely inspect the relationship between the different pathways that were highly correlated with the two dimensional positions and reveal new associations in the data. Considering the top ranked signatures (*p* < 1e – 15 for at least one projection method), and filtering overlapping signatures (Jaccard coefficient > 30%; 63 signatures remaining), we observe a clear pattern of signature clusters (Figure 4). Interestingly, the mesenchymal signature is positively correlated with the expression of coagulation/complement genes (whose expression in the glioblastoma cells studied here was already verified by [2]). Both signatures are also correlated with the TNFα signaling, which supports previous findings concerning the role of this pathway in mesenchymal emergence[24]. On the other hand, the mesenchymal signature is negatively correlated with a hypoxia signature. While hypoxic regions are characteristic in solid tumors[25], this inverse correlation is surprising and possibly aligned with the up-regulation of angiogenetic markers in mesenchymal glioblastoma tumors[22]. Finally, we see a strong negative correlation with a signature of response to the PPAR_*γ*_ agonist rosiglitazone, which aligns with previous observations of beneficial effects PPAR_*γ*_ agonists have in glioblastoma[26, 27, 28]. In addition to the inter-donor variation, FastProject’s visualization also highlights potentially important variation within a tumor. Indeed, the cells from patient MGH31 (Figure 3b, purple) are clearly divided in accordance to the two programs mentioned above - with cells with low mesenchymal and high hypoxic score on one side and the mirror image on the other.

**Figure 4:**
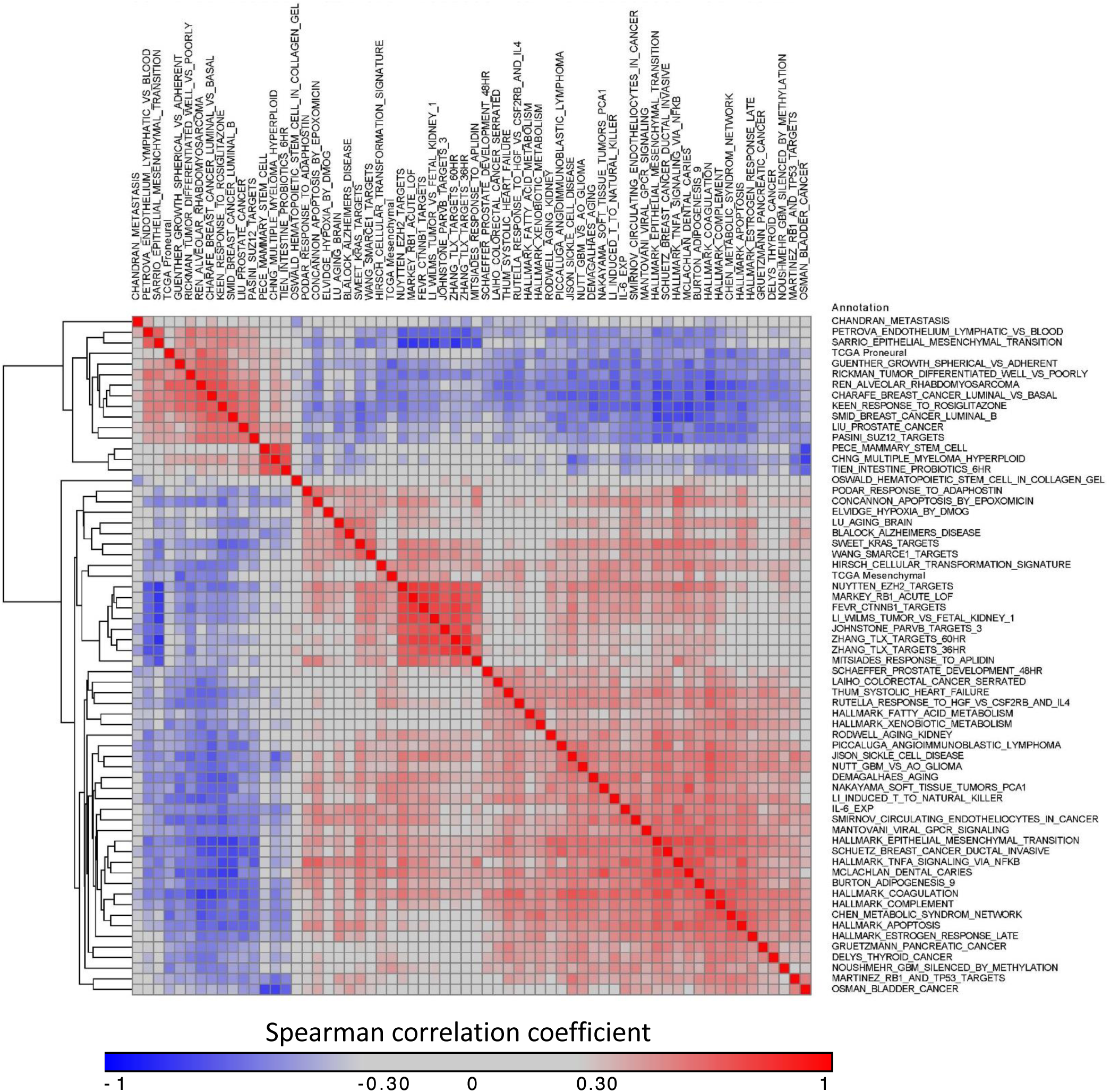
Discovering Correlations between Signatures. FastProject makes its data amenable to further analysis by outputting signature scores and projection coordinates in text format. Shown here is a covariance matrix between top-ranked signatures (*p* < 1e – 15 for at least one projection method) after removing overlapping signatures (*Jaccard coef ficient* > 30%) revealing signatures with similar patterns of expression.

The glioblastoma case study underscores the utility of FastPtroject as a tool for scRNA-Seq data exploration. Starting from a normalized input matrix of gene expression in single cells, and a generic set of signatures, it clearly highlighted some of the main sources for phenotypic variation between cells, and the relation between these sources. Such insights provide an important first step in working with data sets an immense and as complex as the one presented here.

## Conclusions

FastProject is a flexible, comprehensive, and automated pipeline that combines multiple techniques for the analysis of scRNA-Seq. It provides a first glance on the main axes of variation in the data (captured by projections of interest) and their biological meaning (the biological signatures that may explain the organization of cells within the projections). The tool was designed with a flexible API in mind, with the aim of establishing a general platform that will be used by the scRNA-Seq research community for deployment and evaluation of future methods, such as normalization, batch correction, removal of undesired effects (cell cycle, drop-outs), gene/ cell filtering, and dimensionality reduction.

## Methods

### False-Negative estimates

Let *P* (*E*_*ij*_ | *Not D*_*ij*_ ) denote the probability that a gene *i* is expressed in cell *j* conditioned that it is not detected (i.e. probability for a false negative). To estimate this probability we first estimate the priors for the detection, *P*(*D*_*ij*_), and expression, *P*(*E*_*ij*_), events:

For *P*(*D*_*ij*_) we use the percentage of cells that detect gene *i* (*expression* > 0), which we denote as *F*_*i*_. For *P*(*E*_*ij*_), one approach would be to use:

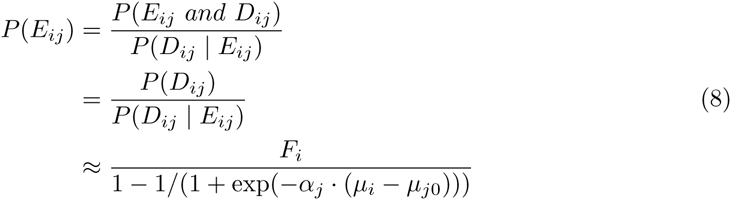

where *μ*_*i*_ is the average of gene *i*’s log expression when detected. However, in order to get a more robust estimation, we use the population mean of the conditional probability,taking: 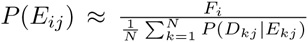 where N is the number of cells in the dataset. For *P*(*Not D*_*ij*_ | *E*_*ij*_) the *F̂*_*j*_(·) functions estimated for each cell (defined in Eq. 1) are used.

Combining these terms, the full expression is:

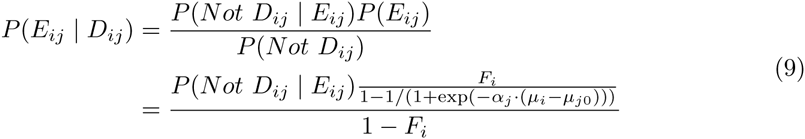

Notably, on occasions where the detection rate in some cells is higher than the prior estimate, this formulation results in a negative value. We therefore restrict the estimate to the range [0,1] by applying a clipping operation.

### Clustering

For each of the projections, the FastProject HTML report includes a simple clustering analysis using the cells’ positioning in the respective two dimensional map. Specifically, samples are clustered based on Euclidean distance in the two-dimensional space using k-means with different *k* values. These clusters are used when rendering a heat-map below the projection showing the (per cluster) average z-score of expression for each gene in the signature.

### Evaluating the consistency of projections and categorical pre-computed signatures

A different method is used to evaluate the significance of signature/projection pairings when operating on pre-computed signatures. For numerical pre-computed signatures, the assigned values are shuffled among cells and for each iteration of this procedure, the signature/projection consistency score is evaluated. P-values are then assessed by comparing the unshuffled score against this distribution using a Z-test as above. For factor signatures, it is necessary to calculated the consistency score in a different manner. To this end, for each pair of signature *S* and projection *p* we compute a signature consistency score representing the extent to which neighboring points in the projection are assigned to the same category. First, we evaluate a neighborhood-consistency score for every cell:

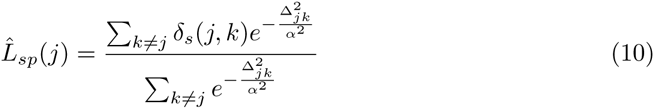

where

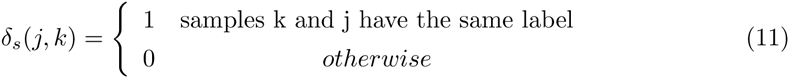

In this way, *L̂*_*sp*_(*i*) is closer to 1 if most of sample *i*’s neighbors have the same label. The signature-projection consistency is then determined by a measure of the overall consistency:

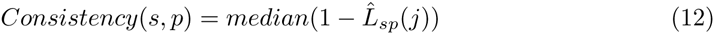

These consistency scores are compared against a distribution of scores calculated from shuffled label assignments to assess significance using a Z-test as above.

## Additional Files

### Additional File 1 — Behavior of Signature Scores: Alternate Data Set

Behavior of signature scores calculated from the LPS-stimulated dendritic cells of Shalek et al 2014 [1]. A) Distribution of Signature/Projection consistency scores across four different types of signatures, Signed (signed immunological signatures from MSigDB), Unsigned (various unsigned hallmark and pathway signatures from MSigDB), Random Signed (signed signatures with randomly selected genes), and Random Unsigned (unsigned signatures with randomly selected genes). B) Distribution of Signature/Projection consistency scores for data in which gene expression levels have been shuffled within each cell. C) Distribution of the Pearson’s correlation coefficient between signature scores and a confounding variable - the proportion of undetected genes in a sample. D) Same as C), but signature scores have been calculated by simply taking the unweighted average of log expression level for genes in the signature. (PNG)

## Additional file 2 — Sample FastProject Output Report

Sample FastProject output html report and associated text files. To view the report, extract the archive and open results.html in a web browser.

## Additional file 3 — Sample FastProject Input Files

Demonstrates how to run FastProject

## Additional file 4 — FastProject software archive

The FastProject package at the time of publication. However, to obtain the most recent version, it is recommended that the reader follows the instructions at the FastProject GitHub repository located at https://github.com/YosefLab/FastProject.

